# Forces and constraints controlling podosome assembly and disassembly

**DOI:** 10.1101/495176

**Authors:** Nisha Mohd Rafiq, Gianluca Grenci, Michael M. Kozlov, Gareth E Jones, Virgile Viasnoff, Alexander D. Bershadsky

**Affiliations:** Mechanobiology Institute, National University of Singapore, Singapore 117411, Singapore; Randall Centre for Cell & Molecular Biophysics, King’s College London, London SE14 5TG, UK; Department of Physiology and Pharmacology, Sackler Faculty of Medicine, Tel Aviv University, 69978 Tel Aviv, Israel; CNRS UMI 3639, 5A engineering drive 1, 117411 Singapore; Department of Biological Sciences, National university of Singapore, 14 Science Drive 4, Singapore 117543; Department of Molecular Cell Biology, Weizmann Institute of Science, Rehovot 76100, Israel

## Abstract

Podosomes are a singular category of integrin-mediated adhesions important in the processes of cell migration, matrix degradation, and cancer cell invasion. Despite a wealth of biochemical studies, the effects of mechanical forces on podosome integrity and dynamics are poorly understood. Here, we show that podosomes are highly sensitive to two groups of physical factors. First, we describe the process of podosome disassembly induced by activation of myosin-IIA filament assembly. Next, we find that podosome integrity and dynamics depends upon membrane tension and can be experimentally perturbed by osmotic swelling and deoxycholate treatment. We have also found that podosomes can be disrupted in a reversible manner by single or cyclic radial stretching of the substratum. We show that disruption of podosomes induced by osmotic swelling is independent of myosin-II filaments. Inhibition of the membrane sculpting protein, dynamin-II, but not clathrin, resulted in activation of myosin-IIA filament formation and disruption of podosomes. The effect of dynamin-II inhibition on podosomes was however independent of myosin-II filaments. Moreover, formation of organized arrays of podosomes in response to microtopographic cues (the ridges with triangular profile) was not accompanied by reorganization of myosin-II filaments. Thus, mechanical elements such as myosin-II filaments and factors affecting membrane tension/sculpting independently modulate podosome formation and dynamics, underlying a versatile response of these adhesion structures to intracellular and extracellular cues.

## Introduction

Podosomes are a distinct type of integrin-mediated cell-matrix adhesion typically seen in cells of monocytic origin (dendritic cells [1, 2], macrophages [3] and osteoclasts [4] but also found more recently in a variety of other cell types [5–8]. In early studies, podosomes were also found in some types of cancer cells (Src-transformed fibroblasts [9, 10]. Thus, podosomes could be considered as an ubiquitous category of integrin-based matrix adhesion structure, whose participation in cancer cell invasion and metastasis is now well documented [11, 12].

Analogous to another class of integrin-based adhesions, focal adhesions, podosomes are membrane specializations in which clusters of integrin family transmembrane receptors are connected to actin filaments. While both integrin receptors themselves and numerous proteins linking the cytoplasmic domains of these receptors with the actin cytoskeleton are similar in podosomes and focal adhesions, the organization of the actin core is different in these types of structures. Focal adhesions are peripheral termini of actin bundles known as stress fibres and the actin scaffold of focal adhesions also consists of bundles of parallel actin filaments [13–15]. Actin filaments of focal adhesions are associated via numerous links containing talin and vinculin with clusters of integrins located immediately underneath the actin filament layer [16]. The major nucleator of actin filaments in focal adhesions are thought to be formins [15, 17–20] even though the Arp2/3 complex plays an important regulatory role especially at the early stage of focal adhesion formation [21, 22]. In contrast, the actin core of podosomes is formed mainly via Arp2/3-driven branching actin polymerization, and Arp2/3 complex along with its activators WASP/N-WASP are essential components of podosomes [3, 23, 24]. In addition, podosomes contain a number of other actin-interacting proteins missing in focal adhesions such as cortactin [25], gelsolin [26], cofilin [27] and dynamin-II [28]. Unlike focal adhesions, the clusters of integrin receptors are located not underneath the actin core but at its periphery forming an approximately ring-shaped structures [29–31]. Formins seem to play less important role in podosome formation than Arp2/3 complex, though the radial filaments connecting the podosome core with proteins of the adhesive ring are postulated to be nucleated by formins [32, 33].

The formation and dynamics of focal adhesions strongly depend on physical forces developed in the course of cell interactions with the extracellular matrix. Treatment of cells with diverse inhibitors interfering with myosin-II filament assembly or mechanochemical activity, as well as knockdown of myosin-IIA, result in disassembly of mature (but not nascent) focal adhesions [34–37]. Moreover, application of external mechanical forces to focal adhesions can promote their growth, while plating on soft or fluid substrates, which do not support development of traction forces by cells, is not favorable for focal adhesion formation [17, 38, 39]. Focal adhesion maturation from initial nascent adhesions as well as their further growth depend on myosin-IIA filament mechanochemical and crosslinking activity as well as on polymerization and bundling of actin filaments[19, 40]. Centripetal traction forces required for focal adhesion growth emerge due to the coupling of centripetal actin flow to integrin clusters via a stick-slip clutch mechanism [41–43]. Formation of myosin-II filament superstructures (stacks)[44] may also play a role in organization of the actin bundles in the lamellum and focal adhesion maturation.

Formation and maintenance of podosomes depends on different mechanical requirements compared to focal adhesions. In particular, podosome formation can efficiently proceed in cells plated on fluid supported membrane bilayers, a substrate which does not permit development of traction forces exerted on integrin clusters [8, 45]. Moreover, fibroblast-type cells, which normally form focal adhesions on stiff matrix, switch to forming podosomes after being plated onto fluid substrates[8]. Thus, podosomes appear to self-assemble by default under conditions of deprivation of traction forces, while focal adhesions critically depend on development of such forces.

In agreement with this premise, recent experimental evidence shows that myosin-II filaments play an inhibitory rather than stimulatory role in podosome formation. Activation of myosin-IIA filament formation by a number of pathways converging to Rho/ROCK signaling axis [46–49] as well as depletion of the regulatory protein S100A4[50] promote podosome disassembly. Proteins known as supervillin and LSP-1 (lymphocyte-specific protein-1) appear to control the disruptive effect of myosin-IIA on podosomes by regulating the recruitment of myosin-IIA to the podosome environs[46, 47].

Last but not least, unlike focal adhesions, which are essentially planar structures, podosomes are known to be small uniform membrane protrusions. This protrusional activity of podosomes is especially evident when cells are attached to soft deformable substrata [8, 51–53]. This means that the processes of membrane sculpting would play an important role in the formation and dynamics of podosomes. Indeed, several proteins known to be potent regulators of membrane sculpting such as dynamin-II[28], and several BAR domain proteins [28, 54, 55] are localized to podosomes and often required for their formation. The effects of mechanical deformation of membrane and factors affecting membrane tension on podosome formation are however not much studied. Furthermore, the relationship between myosin-II driven podosome remodeling and factors affecting membrane sculpting has not been investigated.

The aim of the present study was to explore the effects of different categories of external and internal mechanical forces applied to podosomes. We paid special attention to factors affecting membrane tension as well as processes of membrane sculpting. We show that the effects of factors affecting membrane tension and sculpting on podosomes are independent of the effects of myosin-IIA filaments. We also found that, unlike focal adhesions, podosomes are highly sensitive to substrate stretching and demonstrate unique dependence on substrate topography. Altogether, our results clearly show that podosome formation and dynamics are regulated by two groups of mechanical factors, one operating via myosin-IIA filaments and the other being membrane deformations. This sheds a new light on the unique role of podosome-type adhesions in cell migration, environmental sensing as well as in cancer invasion and metastasis.

## Results

### Disruptive effect of myosin-II filament assembly on podosome integrity

The antagonism between myosin-II filaments overproduction and podosome formation was documented in previous studies[46–50], but the mechanism underlying this antagonism is insufficiently understood. Here we studied the time course of podosome disruption upon activation of myosin-II filament assembly under condition when localization of podosomes is defined by a micropatterned substrate (Figure 1A-D, Supplementary movie 1). We plated THP1 cells differentiated into macrophages by TGFβ1 treatment on micropattened surfaces consisting of 4 μm diameter adhesive islands coated with fibronectin separated by non-adhesive space (Figure 1B). Under such conditions, the podosomes of adherent cells are concentrated in small clusters confined by the islands (Figure 1C). Of note, the intensity of fibronectin fluorescence was significantly lower in zones of the islands underlying podosome clusters, presumably due to podosome-mediated matrix degradation (Figure 1C). The myosin-II filaments in THP1 cells were located at the cell periphery but can also be found in the rims surrounding podosome clusters (Figure 1C). Addition of the RhoA activator, CN03 (Cytoskeleton, Inc; see also ref [56]), triggered myosin-II filament formation within 2 minutes. These filaments are first concentrated at zones surrounding podosome clusters and then move towards the center of the islands occupied by podosomes. The relocation of myosin-II filaments to within 2 μm of podosomes rapidly instigated podosome elimination (Figure 1D and Supplementary Movie 1). Detailed examination of images and movies revealed that the filaments never overlap with existing podosomes. Overall, our data suggests that podosomes disappear as a result of local remodelling of the actin network in their microenvironment induced by myosin-II filaments.

**Figure 1:**
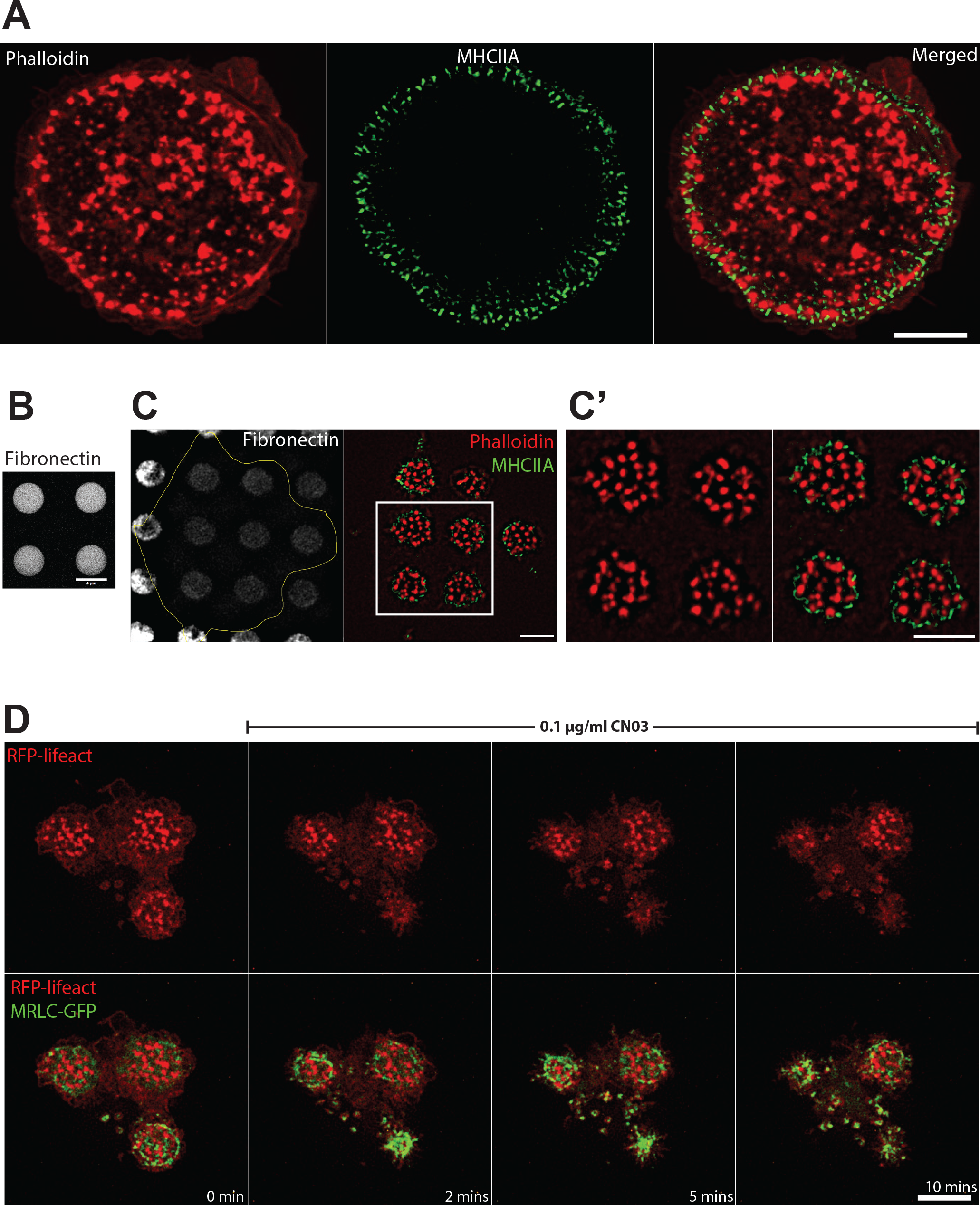
Interrelationship between podosomes and myosin-II filaments in TGFβ1-stimulated THP1 cells. (A) Podosome actin cores visualized by phalloidin staining (red in left panel), and myosin-IIA filaments fluorescently labeled by myosin-IIA heavy chain antibody (green in middle panel); merged image is shown in right panel. Scale bar, 5 μm. Note that myosin-IIA filaments are localized to the narrow peripheral zone of the cell and surround the podosome array. (B) The micropatterned substrate organized as a square lattice consisting of circular fibronectin-coated islands (4 μm diameter) arranged with 8 μm period on passivated non-adhesive substrate. The islands are visualized by fluorescently labeled fibronectin. Scale bar, 4μm. (C) The edge of the cell attached to the fibronectin micropattern (left) is marked by yellow line. Podosomes (actin, red) confined within the adhesive islands are surrounded by myosin-II filaments (green). The boxed area in C is shown at higher magnification in C’, left: actin staining, right: merged image of actin and myosin-IIA filaments. Scale bars, 5 μm. (D) Time course of disruption of podosomes in cells attached to the micropatterned substrate upon activation of RhoA by CN03. The cell attached to three adhesive islands is seen. The cell was transfected with RFP-lifeact (red) and GFP-myosin regulatory light chain (green). Podosomes are localized to the adhesive islands and surrounded by myosin-II filaments. Upper row: actin cores of podosomes, lower row: merged images of podosomes and myosin-II filaments. Note that disruption of podosomes upon addition of CN03 correlates with the assembly of myosin-II filaments. See also Supplementary movie 1. Scale bar, 5 μm.

### Osmotic swelling disperses podosomes and promotes their disassembly

THP1 cells were exposed to RPMI media with 10% serum diluted by water to 50% or 90% dilutions (0.5x or 0.1x hypotonic, respectively) for 15 minutes. Such treatments are known to increase membrane tension[57–60]. Cells incubated with 0.5x hypotonic medium showed reductions in podosome number as compared to cells treated with isotonic medium (Figure 2A and B, graphs 2D and E). The sparse residual podosomes were approximately 5-fold smaller in area than podosomes in control cells (Figure 2B and F, Supplementary Figure 1A, C-D and Supplementary Movie 2) as revealed by structured-illumination microscopy (SIM). The cells demonstrated numerous lamellipodia and increased spread area (Figure 2F). The effect of 0.5x hypotonic medium on podosomes was transient, so that new podosomes appeared within an hour following medium dilution. Incubation in 0.1x hypotonic medium (90% dilution) resulted in cell retraction and formation of numerous irregular actin-rich protrusions (Figure 2C). Such cells do not demonstrate any podosome-like structures for several hours. Since we demonstrated above that podosome disruption is induced by activation of myosin-II filament formation, we examined whether elimination of myosin-II filaments interfered with the effect of hypo-osmotic shock. We found that in spite of complete disassembly of myosin-II filaments upon treatment with 30 μM Rho kinase inhibitor Y-27632, the 0.5x hypotonic medium still disrupts podosomes (Figure 2G, Supplementary movie 3). Unlike podosomes, focal adhesions of mouse embryonic fibroblast (MEFs) did not disassemble upon hypoosmotic shock (Figure 2H, Supplementary movie 4).

**Figure 2:**
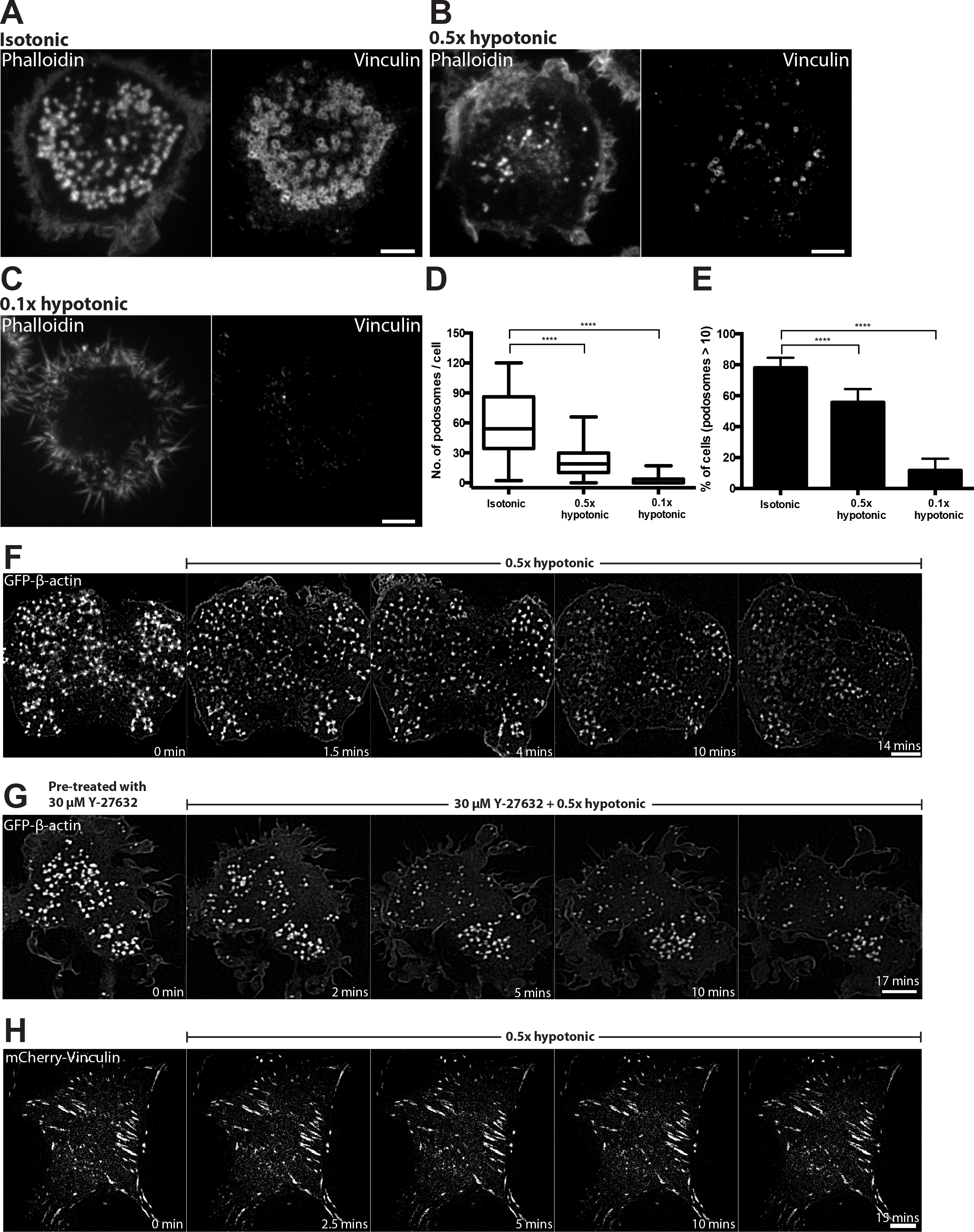
Disruption of podosomes upon osmotic swelling. (A) Cells exposed for 15 minutes to isotonic (complete medium) and (B and C) hypotonic medium with 50% (B) and 90% (C) reduction in osmolarity. F-Actin (left) and vinculin (right) were visualized by phalloidin and vinculin antibody staining, respectively. Scale bars, 5 μm. (D and E) Graphs showing the number of podosomes per cell presented as box-and-whiskers plots (D) and the percentage of cells with more than 10 podosomes presented as mean ± SD (E). In this and the following figures, the significance of the difference between groups was estimated by two-tailed Student’s t-test, and the range of p-values >0.05(non-significant), ≤ 0.05, ≤0.01, ≤0.001, ≤ 0.0001 are denoted by “ns”, one, two, three and four asterisks (*), respectively. (F) Time-lapse structured-illumination microscopy (SIM) visualization of podosome dynamics in GFP-β-actin-transfected cell exposed to 0.5x hypotonic medium. Note dispersion and reduction in the number of podosomes upon hypotonic shock. See Supplementary movie 2. (G) Rho kinase inhibitor, Y-27632, did not prevent the disruption of podosomes upon hypotonic shock. Scale bars, 5 μm. See Supplementary movie 3. (H) Focal adhesions of mouse fibroblasts visualized by expression of mCherry-vinculin were not affected by incubation in 0.5x hypotonic medium. Scale bar, 10 μm. See Supplementary movie 4.

### Decreasing membrane tension by deoxycholate promoted the clustering of podosomes

In order to decrease membrane tension without perturbing the spread area of the cell, small concentrations of the detergent, deoxycholate, were used to expand the lipid bilayer[59]. Addition of 400 μM deoxycholate induced immediate disruption of podosomes, which however quickly recovered with the formation of large podosome clusters (Figure 3A, B, F, Supplementary movie 5). As a result, by 5 minutes following deoxycholate addition, podosome number was similar to that in non-treated cells but their organization differed significantly. Instead of individual podosomes evenly distributed over the cell ventral surface, podosomes in deoxycholate-treated cells were organized into clusters containing more than 50 podosomes each (graphs 3C and D). Visualization of podosomes with high magnification using SIM, revealed in control cells thin actin links connecting neighbouring podosomes. Such links were significantly shortened or entirely disappeared in the podosome clusters formed in deoxycholate-treated cells (Figure 3B inset, graph 3E). The average distance between individual podosomes in the cluster was 0.24 ± 0.02 μm compared with 0.54 ± 0.04 μm in controls, so podosomes were tightly-packed (graph 3E). There were no differences in actin core diameter or actin fluorescence intensity between podosomes in deoxycholate-treated and control cells (Supplementary Figure 1B, C-D). The clustering effect induced by deoxycholate was however transient and podosomes return to a normal distribution at about 30 minutes after addition of deoxycholate (Figure 2G, Supplementary movie 6).

**Figure 3:**
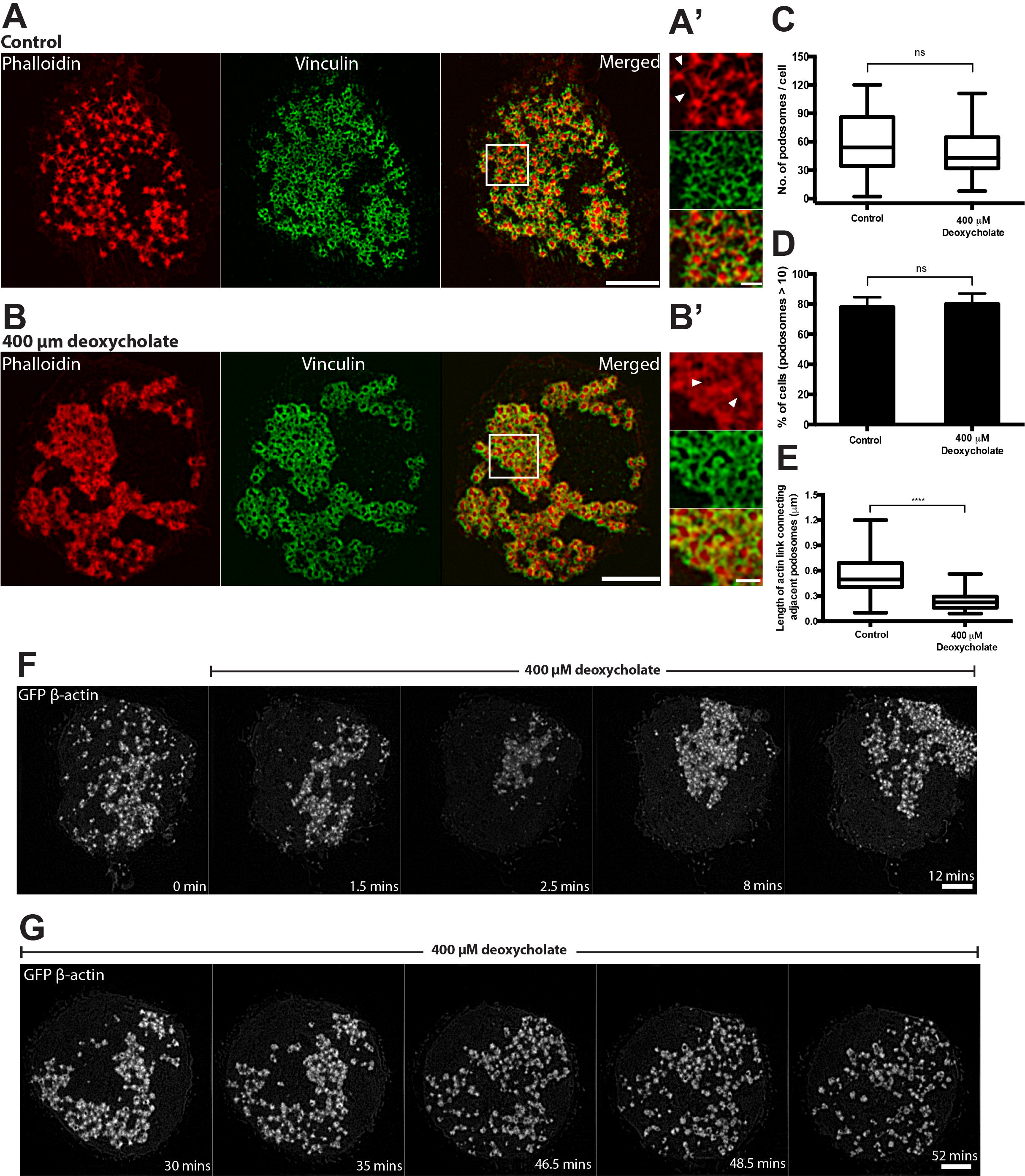
Decreasing membrane tension by the detergent deoxycholate promotes podosome cluster formation. Structured-illumination microscopy (SIM) visualization of podosomes in THP1 cells exposed to (A) normal complete medium and (B) 400 μM deoxycholate-containing medium for 15 minutes. Actin labeled with phalloidin (left, red), vinculin visualized by antibody staining (middle, green) and their merged image (right) are shown. Scale bars, 5 μm. Boxed areas in A and B are shown at higher magnification in A’ and B’, respectively. Scale bars, 1 μm. White arrowheads showing actin links between neighboring podosomes. (C-E) Quantification of 15 minutes-deoxycholate treatment on podosomes. (C) Number of podosomes per cell presented as box-and-whiskers plots. (D) Percentage of cells with more than 10 podosomes presented as mean ± SD. (E) Length of the actin links between podosomes. The p-values are marked by asterisks as described in the legend to Figure 1. (F and G) Time-lapse images showing the evolution of podosomes upon addition of deoxycholate. (F) First 30 minutes following deoxycholate addition. See Supplementary Movies 5. (G) 30-60 minutes after addition of deoxcycholate. See Supplementary Movies 6. The cells were transfected with GFP-β-actin to visualize the actin cores of podosomes. Scale bars, 5 μm. Note severe decrease of podosome number at 1.5-2 minutes following deoxycholate addition, formation of new podosomes organized into clusters at 8-35 minutes of incubation with deoxycholate and recovery of uniform podosome distribution at 35 minutes onwards.

### Inhibition of dynamin-II but not clathrin-mediated endocytosis perturbed podosome formation

Since physical factors increasing membrane tension are known to inhibit endocytosis[58, 61–64], we considered whether processes of endocytosis are involved in podosome formation and maintenance. First, we confirmed and extended previous data on the role of dynamin-II in podosomes [28] showing that inhibition of dynamin-II either by pharmacological inhibitor, dynasore, or by knockdown of dynamin-II, resulted in severe disruption of podosomes in THP1 cells and the formation of focal adhesions instead (Figure 4A-E). Inspection of myosin-II organization in cells with dynamin-II inhibition revealed that such cells have significantly increased numbers of myosin-II filaments (Figure 4C) as compared to functional dynamin-II-containing THP1 cells. These myosin-II filaments occupied the entire cytoplasm of dynasore-treated or dynamin-II knockdown cells but did not form regular myosin-II filament stacks (Figure 4C). Since we have shown above that accumulation of myosin-II filaments leads on its own to podosome disruption, we examined whether accumulation of myosin-II filaments participate in the podosome disruption induced by dynamin-II inhibition. When dynasore was added to THP1 cells pre-treated with ROCK inhibitor Y-27632 for 30 minutes, podosomes were still disrupted in spite of the complete absence of myosin-II filaments in these cells (Figure 4D). Of note, focal adhesions typical for dynasore-treated cells did not form in cells pre-treated with Y-27632 (Figure 4C and D). Similarly, treatment with Y-27632 was insufficient to rescue podosomes in dynamin-II knockdown THP1 cells (Figure 4E). These data strongly suggests that accumulation of myosin-II filaments did not participate in the disruption of podosomes induced by dynamin-II suppression. In addition, while dynamin inhibition showed a severe disruptive effect on podosomes, the inhibitor of clathrin-mediated endocytosis, Pitstop^®^2, did not interfere with podosome integrity in our experiments (Figure 4E).

**Figure 4:**
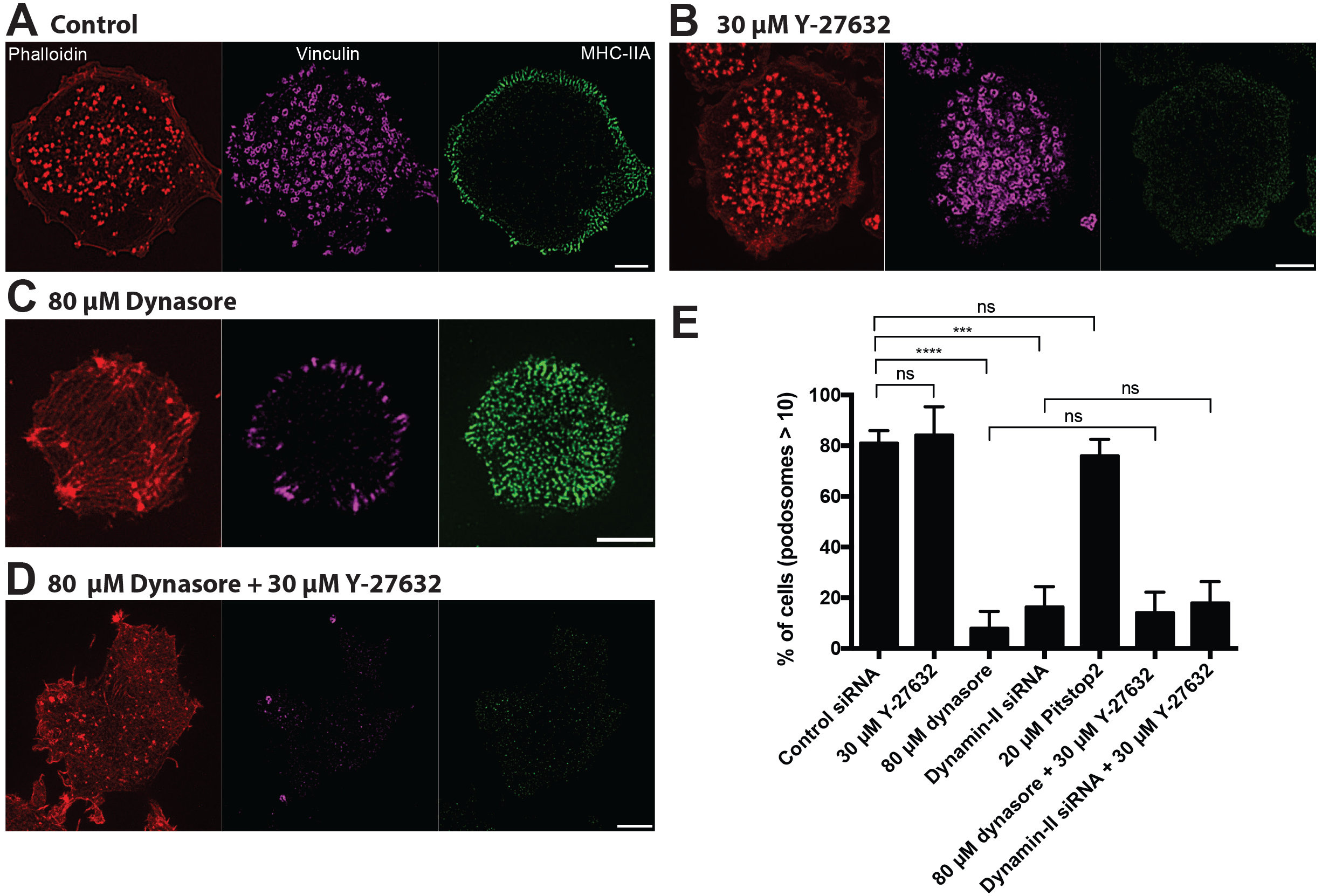
Dynamin-II inhibition or knockdown disrupts podosomes in a myosin-II independent manner. Images of cells treated with (A) 0.1% DMSO, (B) 30 μM Y-27632, (C) 80 μM dynasore, (D) 80 μM dynasore added 30 minutes after addition of 30 μM Y-27632 are shown. The cells were fixed 1 hour after addition of corresponding drugs. Actin podosome cores were visualized by phalloidin (left, red); podosome rings (middle, purple) and myosin-IIA filaments (right, green) were visualized by immunofluorescence staining of vinculin and myosin-IIA heavy chain. Scale bars, 5 μm. Note that dynasore treatment disrupted podosomes and induces formation of new myosin-II filaments (C), but inhibition of myosin-II filaments formation by Y-27632 (B) did not prevent the disruptive effect of dynasore on podosomes (D). Quantification of the percentage of cells having more than 10 podosomes for the treatments shown in A-D, as well as for dynamin-II siRNA knockdown cells treated or untreated with Y-27632, and cells treated with the clathrin inhibitor Pitstop®2. Note that dynamin-II knockdown prevented podosome formation irrespectively to treatment with Y-27632, which disrupts myosin-II filaments. Clathrin inhibitor Pitstop®2 did not affect podosome integrity. The significance of the differences between percentage of podosome-forming cells after different treatments (p-values) are marked by asterisks as described in the legend to Figure 1.

### Radial stretching of the substrate induces loss of podosomes

Membrane tension can also be augmented by physical stretching of the substrate to which the cell is attached [58, 65]. In our study, differentiated THP1 cells were plated on PDMS surfaces coated with fibronectin which were stretched radially using a previously described device [65].

A 5% single radial stretch maintained for 10 seconds as well as a 5% cyclic stretch (0.1Hz, 6 hours) resulted in a modest but significant decrease in podosome number per cell (Figure 5A and D, Graph F) while the percentage of podosome-containing cells did not decrease after such treatment (Graph 5G). When the stretch magnitude was increased to 15%, podosomes were completely disrupted (Figure 5B and E). During such stretch, the cells formed numerous blebs (Figure 5B and E). In spite of total podosome disassembly, cells remained attached to the substrate. The podosomes did not recover until the stretch was released; following stretch release, both the re-assembly of podosomes and the disappearance of the blebs was seen, with complete recovery of the control phenotype in 30 minutes following the release of stretch (Figure 5C). Cycles of substrate stretching and release can be repeated several times with reproducible reverse responses of podosomes and blebs. The results were not affected if the duration of single stretch was increased up to 1 minute, and cyclic stretch frequency varied from 0.01 to 0.1 Hz. Collectively, the data show that podosomes respond strongly to the magnitude of stretch but not to the duration of a single stretch or frequency of cyclic stretch, and cells are capable of forming new podosomes once the strain was released (Graphs 5F and G).

**Figure 5:**
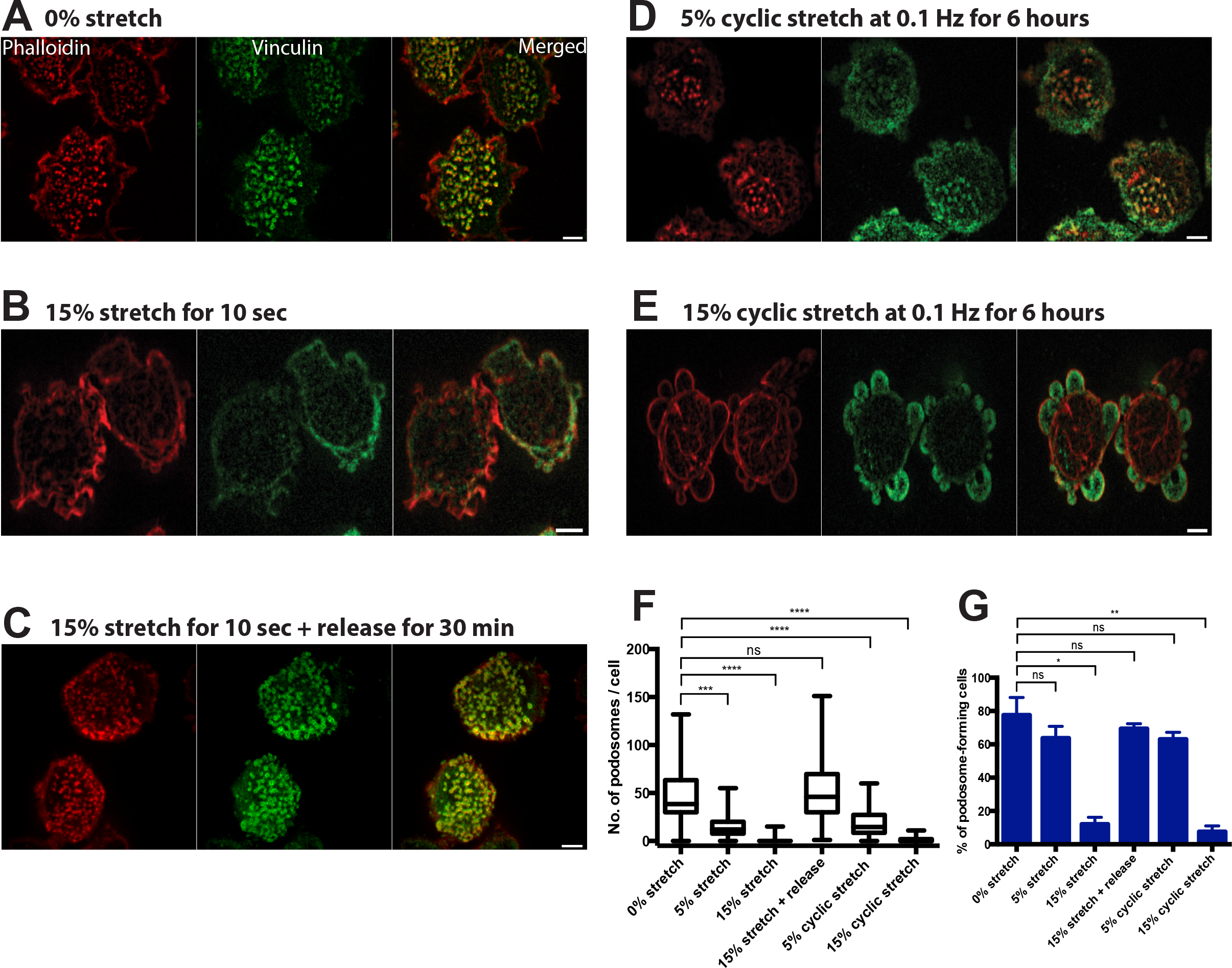
Stretching of substrate induced podosome disassembly. (A-E) Representative images of cells on PDMS (polydimethylsiloxane) substrate coated with fibronectin fixed and stained after substrate radial stretching or release. Actin was labeled with phalloidin (left, red), vinculin was visualized by antibody staining (middle, green) and their merged image is shown in the right panel. (A) Cells on non-stretched substrate. (B) Cells subjected to 15% stretching for 10 seconds. (C) Cells stretched for 10 seconds and incubated for 30minutes following the stretch release. (D and E) Cells subjected to cyclic stretching of (D) 5% and (E) 15% magnitude at 0.1 Hz for 6 hours. Scale bars, 5 μm. (F and G) Quantification of (F) the number of podosomes per cell (box-and-whiskers plots), and (G) percentage of cells containing more than 10 podosomes (mean ± SD) for all experimental situations shown in (A-E), as well as for 5% single stretch for 10 seconds. Graphs represent results of 2-3 independent experiments.

### Formation of linear podosome arrays induced by substrate topography

Another group of physical factors affecting podosome formation are topographical features of the substrate. It was shown previously that podosomes of dendritic cells prefer to form in the proximity of 90° steps on the substrate[66] or can be induced by pores in the nuclearpore™ filters[67]. To further explore the effects of 3D topography on podosome organization, THP1 cells were plated on arrays of microfabricated ridges with triangular profile of various heights (Figure 6A) arranged with different pitches (Figure 6B). We have found that cells plated on fibronectin-coated substrates with such ridges showed highly ordered podosome formation, which formed double chains along each side of the base of the triangular ridge regardless of their heights or the spacing between them (Figure 6C-F). Cells were capable of forming single or multiple streaks of double chains of podosomes depending on the spacing between the ridges (Figure 6C-F). Furthermore, these podosomes preferred the sides of the crest to the flat area of the substrate, suggesting that sensing local changes in topography can favor the assembly of podosomes in an organized linear fashion.

**Figure 6:**
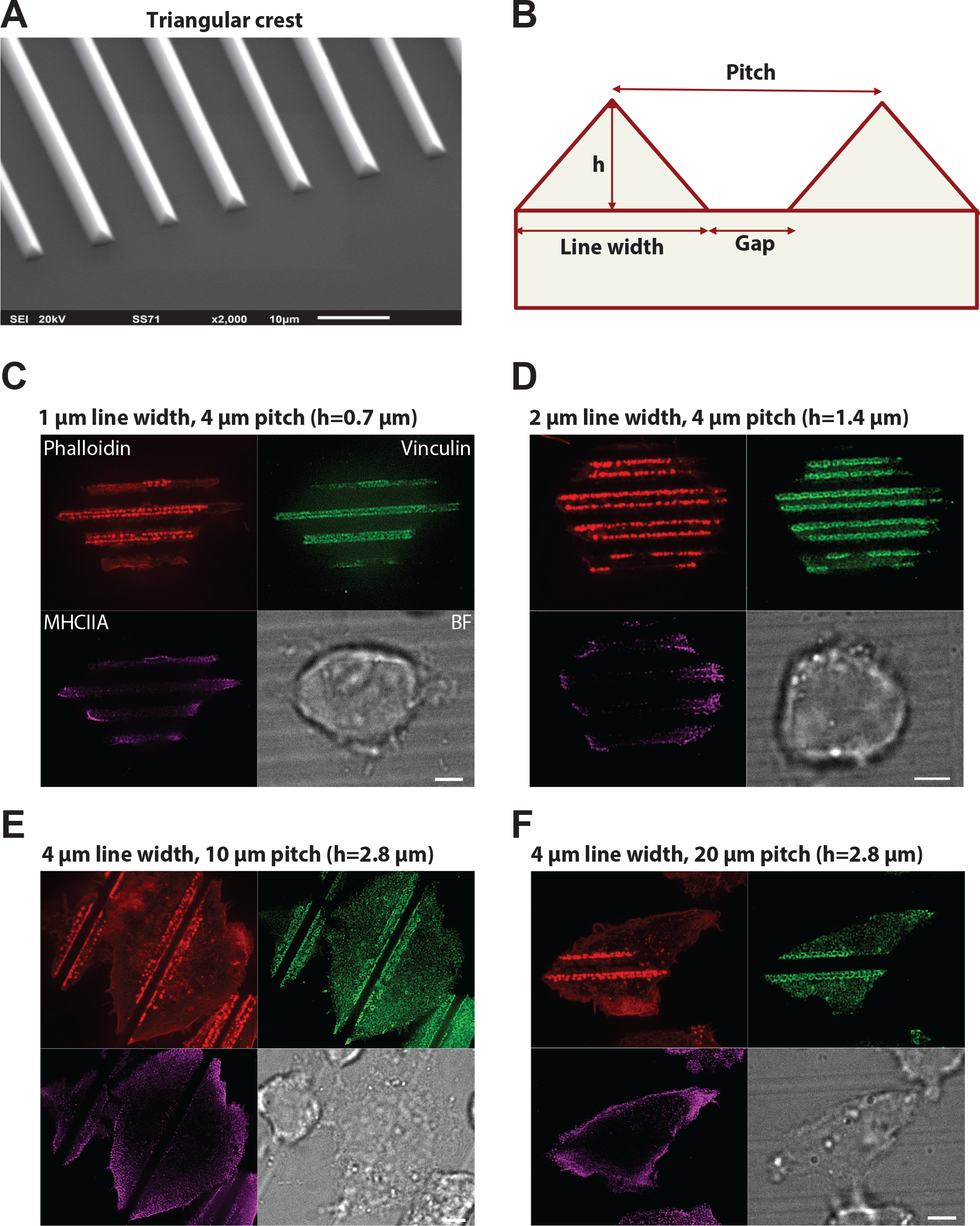
Podosome formation in response to topographical cues. (A) Scanning electron microscopy image of PDMS substrate with triangular ridges. (B) Schematic diagram of profile of the substrate with triangular ridges. The PDMS substrate was coated with fibronectin (1 μg/ml) for 1 hour. The width of the ridges, distances between crests (pitch) and ridges’ height (h) were varied as shown in (C-F). In each panel, four views of the same cell are presented: actin cores visualized by phalloidin staining (upper left, red), podosome adhesive domains visualized by vinculin staining (upper right, green), distribution of myosin-IIA filaments (lower left, purple), and bright field image of the cell (lower right). Scale bars, 5 μm. Note that in all cases podosomes are aligned along both sides of the ridges, and not formed in other regions of the cell. At the substrate with smallest ridges (pitch: 1 μm), the discrete actin cores are seen but vinculin is not organized into ring-shaped structures. At pitches 2-20 μm, chains of typical podosomes are seen. Myosin-IIA filaments are located at the cell periphery similarly to the case of cells plated on planar substrate.

Examination of distribution of myosin-IIA filaments in cells on the patterned substrates revealed that filaments remained at the cell periphery, forming a sub-cortical ring similar to that seen in cells on a flat substrate (Figure 6C). This suggests that formation of podosome arrays at the edges of the triangular crests as well as disappearance of podosome from the other regions of the cell is not a consequence of reorganization of myosin-IIA filaments.

## Discussion

In this study, we explored the effects of several types of factors on the integrity of podosomes in macrophage-type cells. We confirmed and extended the disruptive effect of myosin-IIA filaments on podosomes. By direct observation of the interaction of small groups of podosomes with surrounding myosin-IIA filaments, we found that activation of myosin-II filament formation affects podosomes locally, and the distance at which myosin-II filaments exert their effect is in the micron range. The process of podosome disassembly upon local activation of myosin-II filament formation was rapid and lasted less than a minute. The disassembly of myosin-II filaments, known to have a profound disruptive effect on focal adhesion integrity[17, 34–37], does not affect podosomes.

There are several not necessary mutually exclusive mechanisms by which excessive assembly of myosin-II filaments can interfere with podosome integrity. Myosin-II filaments under certain conditions can depolymerize actin [68, 69] or disintegrate branching actin networks nucleated by Arp2/3 complex. Forces generated through interaction of myosin-II and actin filaments may therefore interfere with the Arp2/3-mediated actin polymerization in podosomes. Finally, myosin-II filaments underlying the plasma membrane could affect membrane tension or curvature [70, 71] in such a way that hinders formation of podosomal protrusions.

We therefore investigated the role of factors affecting membrane tension and sculpting on podosome integrity and dynamics. We have clearly shown that hypoosmotic shock known to increase membrane tension results in disassembly of podosomes. Of note, suppression of myosin-II filament formation by Rho kinase inhibitior Y-27632 does not prevent or rescue the disruptive effect of hypo-osmotic shock on podosomes. The decrease of membrane tension by treatment with deoxycholate also exerts profound effect on podosomes resulting in podosome clustering.

Dynamin-II, known for its function in membrane fission[72] and regulation of Arp2/3 complex-driven actin polymerization, is required for podosome integrity [28, 73]. We checked whether the disruptive effect of dynamin-II inhibition on podosomes is mediated by myosin-IIA filament formation. This conjecture was motivated by our observation that treatment of THP1 cells with dynamin inhibitor, Dynasore, triggered formation of multiple myosin-IIA filaments. However, inhibition of myosin-IIA filament formation by Rho kinase inhibitor did not prevent the disruptive effect of dynamin inhibition on podosomes. Thus, not all podosome-disrupting factors operate via myosin-II filaments and in particular manipulations with membrane tension or sculpting can affect podosome directly.

We further investigated the effects of single or cyclic substrate stretching on podosome integrity and clearly showed that such manipulations result in reversible disassembly of podosomes. This effect can be explained by the increase of membrane tension upon stretching observed in previous studies[58, 74]. However, in addition to membrane tension, the substrate stretching was shown to activate RhoA[75] and reinforce contractility[76] which suggests that formation of myosin-IIA filaments is activated. Thus, effect of substrate stretching on podosomes is most probably a result of combined action of increased membrane tension and augmented myosin-II filament formation.

A common feature of all podosome-disrupting treatments discussed above is their inhibitory effect on endocytosis. Indeed, it is well-documented that increase in membrane tension via osmotic swelling inhibits the endocytosis[58, 61–64]. Substrate stretching is also known to interfere with endocytosis[58]. Finally, dynamin, a protein required for multiple forms of endocytosis [77] is indispensible for podosome integrity. The sensitivity of podosomes to factors inhibiting endocytosis can be explained by the possible role of endocytosis in the membrane balance at podosomes. Podosomes seem to be the sites of intense insertion of new membrane required for both protrusional activity and exocytosis of vesicles containing metalloproteinases degrading the extracellular matrix [11]. Therefore, compensatory endocytosis[78] could be needed for the maintenance of membrane equilibrium. Since inhibition of clathrin-mediated endocytosis did not affect podosome integrity in our experiments, the endocytosis that might be required for podosome integrity is probably clathrin-independent.

Of note, the sensitivity of both endocytosis and podosome formation to similar experimental treatments can also reflect the similarity between the processes of endocytosis and podosome formation. In both cases, the mechanism depends on membrane remodeling mediated by the actin cytoskeleton. Further elucidation of this analogy is an interesting avenue for future studies.

The unique dependence of podosome formation on substrate topography described in this study and previous publications [66, 67] provides another example of podosome regulation by membrane remodeling. Indeed, we have found that preferential formation of podosomes along the triangular ridges of microfabricated substrates is not accompanied by apparent changes in the distribution of myosin-II filaments. Thus, the factors inducing podosome alignment along ridges and suppressing podosome formation in other cell regions depend rather on the shaping of the membrane and submembrane cortex. Further studies are needed to elucidate the mechanism of such membrane shaping-dependent regulation of podosomes.

Summing up, our study revealed two groups of mechanisms affecting podosome integrity and dynamics involving myosin-IIA filament reorganization and membrane tension/shape changes, respectively. These mechanisms seem to be operating for podosomes rather than other types of integrin adhesions, such as focal adhesions. Further elucidation of these mechanisms will be important for understanding of podosome-dependent processes where these are of important biological consequences such as 3D cell migration and tumor cell invasion.

## Acknowledgements

This research is supported by the National Research Foundation, Prime Minister’s Office, Singapore and the Ministry of Education under the Research Centres of Excellence programme (A.D.B, N.B.M.R., G.G., V.V.) and Singapore Ministry of Education Academic Research Fund Tier 3 (A.D.B, V.V.) MOE Grant No. MOE2016-T3-1-002). N.B.M.R is also funded by a joint National University of Singapore-King’s College London graduate studentship. G.E.J. is supported by the Medical Research Council, UK (G1100041, MR/K015664) and the generous provision of a visiting professorship from the Mechanobiology Institute, Singapore.

## Materials and Method

### Cell culture, plasmid and transfection procedures

THP1 human monocytic leukemia cell line was obtained from Health Protection Agency Culture Collections (Porton Down, Salisbury, UK) and cultured using Roswell Park Memorial Institute (RPMI-1640) supplemented with 10% HI-FBS and 50 μg/ml 2-Mercaptoethanol (Sigma-Aldrich) at 37°C and 5% CO_2_.

The suspension-cultured THP-1 cells were differentiated into adherent macrophage-like cells either with 1 ng/ml human recombinant cytokine TGFβ1 (R&D Systems) for 24 or 48 hours on fibronectin-coated glass substrates. No apparent difference between the phenotype of cells stimulated for 24 or 48 hours were detected. 35-mm ibidi (Cat. 81158) glass-bottomed dishes were coated with 1 μg/ml of fibronectin (Calbiochem, Merck Millipore) in phosphate buffered saline (PBS) for at least 1 hour at 37°C, washed with PBS twice, and incubated in complete medium containing TGFβ1 prior to seeding of cells.

THP1 stably expressing GFP-β-actin (described in Cox, Rosten [79]), or stably co-expressing RFP-lifeact and human GFP-myosin regulatory light chain (MRLC) (described in Rafiq, Lieu [48], Rafiq, Nishimura [80]) were used in all live experiments concerning podosome studies.

For dynamin-II knockdown, THP1 cells were transfected with 100nM of dynamin-II siRNA (Dharmacon, ON-TARGETplus SMARTpool siRNA, catalogue no. L-004007-00-0005). For control experiments, non-targeting pool siRNA (Dharmacon, ON-TARGETplus, catalogue no. D-001810-10) was used at a similar concentration. Cells were transfected using electroporation (Neon Transfection System, Life Technologies) in accordance to manufacturer’s instructions. Specifically, two pulses of 1400V for 20 milliseconds were used.

Immortalized rptp-α(+/+) mouse embryonic fibroblasts (Su et al 1999) that was termed MEFs, were obtained from the Sheetz laboratory (Mechanobiology Institute, Singapore). MEFs were cultured in Dulbecco’s modified Eagle’s Medium high glucose (DMEM), supplemented with 10% heat-inactivated fetal bovine serum (HI-FBS, Gibco), 1% L-Glutamine, and 100 IU/mg penicillin-streptomycin (Invitrogen) at 37°C and 5% CO_2_. MEFs were either seeded on fibronectin-coated 35-mm ibidi or 27-mm IWAKI (Japan) glass-bottomed dishes for 24 hours post-transfection.

For focal adhesion analysis, MEFs were transiently electroporated (Neon Transfection System, Life Technologies) with mCherry- Vinculin (Dr. Michael W. Davidson, Florida State University, FL, USA) with a single pulse of 1400V for 20 milliseconds.

### Immunofluorescence

THP1 Cells were fixed for 15 minutes with 3.7% PFA in PBS, washed twice in PBS, permeabilized for 10 minutes with 0.5% triton X-100 (Sigma-Aldrich) in PBS, and then washed twice again in PBS. For microtubule visualization, cells were fixed and simultaneously permeabilized for 15 min at 37°C in a mixture of 3% PFA–PBS, 0.25% Triton-X-100 and 0.2% glutaraldehyde in PBS, and then washed twice with PBS for 10 min. Before immunostaining, samples were quenched for 15 min on ice with 1 mg/ml sodium borohydride in cytoskeleton buffer (10 mM MES, 150 mM NaCl, 5 mM EGTA, 5 mM MgCl_2_, 5 mM glucose, pH 6.1). Fixed cells were blocked with 5% BSA or 5% FBS for 1 hour at room temperature prior to incubation with the following primary antibodies overnight at 4°C: anti-tubulin (Sigma-Aldrich, catalogue no. T6199, dilution 1:400); anti-vinculin (Sigma-Aldrich, catalogue no. V9131, dilution 1:400); anti-non muscle heavy chain of myosin-IIA (Sigma-Aldrich, catalogue no. M8064, dilution 1:500); Samples were washed with PBS three times and incubated with Alexa Fluor-conjugated secondary antibodies (Thermo Fisher Scientific) for 1 hour at room temperature, followed by three washes in PBS. F-actin was visualized by Alexa Fluor 488 Phalloidin (Thermo Fisher Scientific), Phalloidin-TRITC (Sigma-Aldrich) or Alexa Fluor 647 Phalloidin (Thermo Fisher Scientific).

### Osmotic shock and drug treatments

Pharmacological treatments were performed using the following concentrations of inhibitors or activators: 30 μM for Y-27632 dihydrochloride (Sigma-Aldrich), 80 μM for dynasore (Sigma-Aldrich), 400 μM deoxycholate (Sigma-Aldrich), and 0.1 μg/ml for Rho Activator II (CN03, Cytoskeleton). Duration of the treatment with the inhibitors was 1 hour unless otherwise stated. In some cases, cells were pre-treated with one inhibitor for 30 min and then another inhibitor was added for additional 1 hour. For hypotonic experiments, cells were exposed to solutions containing complete medium diluted in sterile water at 1:1 (0.5x hypotonic) or 1:9 (0.1x hypotonic) using a perfusion chamber (CM-B25-1, Chamlide CMB chamber) and image was acquired just prior to the start of acquisition.

### Micro-patterning of adhesive islands using UV-induced molecular adsorption

Adhesive islands with a diameter of 4 μm were printed in square lattices with a period of 8 μm. Clean glass coverslips were sealed with NOA 73 liquid adhesive to plastic-containing dishes by UV treatment for 2 minutes, and were then treated with oxygen plasma for 5 minutes. The coverslips were coated with PLL-g-PEG (PLL(20)-g[3.5]-PEG(5), SuSoS AG, Dübendorf, Switzerland) at 100 μg/mL in PBS for at least 8 hours followed by multiple washes with PBS. For micropattern printing, PRIMO system (Alveole, France) mounted on an inverted microscope (Nikon Eclipse Ti-E, Japan) equipped with a motorized scanning stage (Physik Instrumente, Germany) was used as Digital Micro-mirror Device (DMD) to create a UV pattern at 105 μm above the focal plane of the microscope. After alignment of specimen with the UV pattern by using the Leonardo software (Alveole, France), a solution of photoinitiator (PLPP, Alveole, France) was incubated on the dishes. Depending on the UV exposure and time, the UV-activated photoinitiator molecules locally cleaved the PEG chains, which permits subsequent local deposition of proteins. The dishes were washed with PBS multiple times prior to incubation with labeled fibronectin (Alexa 488 Fibronectin, Cytoskeleton, Inc., 50 μg/ml) or unlabeled fibronectin mixed with Fibrinogen Alexa 647 (ThermoFisher Scientific, 50 μg/mL) at a ratio of 20:1 for 10 minutes. After several washes with PBS, THP1 cells were plated on these micropatterned lattices for SIM imaging.

### Cell stretching assay

THP1 cells were subjected to 0, 5 or 15% single or cyclic radial stretch using the stretching device described in [65]. Briefly, cells were plated on a layer of polydimethylsiloxane (PDMS) coated with 10 μg/ml fibronectin, in a stretching unit. The substrate stretching was generated via changing the pressure in a chamber underneath the stretchable substrate. For single stretch experiments, cells were incubated under stretched conditions for 10 seconds, and then fixed as described above. The stretching itself lasted for less than a second[65]. For single stretch recovery experiments, cells were released from stretching 30 minutes prior to fixing. For cyclic stretching, cells were exposed to stretching with a frequency of 0.1 Hz at 5 or 15% stretch magnitude and then fixed.

### Fluorescence microscopy

THP1 cells (in Figure 2A-C, Figure 5) and MEFs (in Figure 2H) were imaged in complete medium (unless stated otherwise) using a spinning-disc confocal microscope (PerkinElmer Ultraview VoX) attached to an Olympus IX81 inverted microscope, equipped with a 100x oil immersion objective (1.40 NA, UPlanSApo) and EMCCD camera (C9100-13, Hamamatsu Photonics) for image acquisition. Volocity software (PerkinElmer) was used to control the image acquisition. For all other images, two types of structured illumination microscopy equipments were used: 1) spinning-disc confocal microscopy (Roper Scientific) coupled with the Live SR module [81], Nikon Eclipse Ti-E inverted microscope with Perfect Focus System, controlled by MetaMorph software (Molecular device) supplemented with a 100x oil 1.45 NA CFI Plan Apo Lambda oil immersion objective and sCMOS camera (Prime 95B, Photometrics), 2) Nikon N-SIM microscope, based on a Nikon Ti-E inverted microscope with Perfect Focus System controlled by Nikon NIS-Elements AR software supplemented with a 100x oil immersion objective (1.40 NA, CFI Plan-ApochromatVC) and EMCCD camera (Andor Ixon DU-897).

### Microfabrication of triangular ridges

The coverslips with PDMS structures on top have been produced adapting a molding protocol from Ref [82]. Silicon molds 15×15 mm^2^ wide with 1.5 mm long trenches of triangular cross section with different sizes were prepared *via* silicon anisotropic etching. Briefly, standard silicon wafers with 300 nm of SiO_2_ thermally grown on both side were spin-coated with 1 μm thick AZ214E positive tone photo-resist. The pattern was then produced with direct writing in a DWL-66fs Heidelberg laser writer equipped with a diode-laser at 375 nm. After development for 1 min in AZ400K diluted 1:4 in DI water, the patterned resist mask is then used to etch the silicon oxide layer in a Samco 10NR RIE tool using CF_4_/O_2_ etching chemistry (40/4 sccm respectively, 15 Pa, 150 W applied through an RF generator at 13.56 MHz, as described in [83]. After stripping the resist, 10 min of anisotropic etching in 5 M KOH at 80 °C produced the triangular trenches with the designed sizes. After the anisotropic etching, the silicon oxide is removed with immersion in a buffered oxide etching (a solution of 1:7 of HF: NH_4_F in water, this etching solution is selective for silicon oxide but does not attack Si). The wafer is then diced in the 16 single dyes, and each is coated with an anti-sticking self-assembled monolayer of Trichloro(1H,1H,2H,2H-perfluorooctyl)silane by vapour deposition. PDMS (Sylgard 184, Dow Cornig, USA) is prepared in 10:1 ratio with its reticulation agent and degassed for 30 min in a vacuum jar after careful mixing. A 10 μm layer is spin-coated on the coverslip (4000 rpm for 60 s) and degassed a second time for 10 min. A silanized mold is then placed on top of the PDMS coating and gently pressed to the coverslip with around 150 kPa of pressure to facilitate the filling of the triangular cavities. While keeping the pressure applied, the assembly is transferred on a Hot Plate and the temperature risen to 120 °C, where PDMS reticulation is left proceeding for 30 min. After cooling down to room temperature, the silicon mold is peeled off revealing the structured PDMS layer on top of the coverslip with the triangular ridges.

**Supplementary Figure 1:**
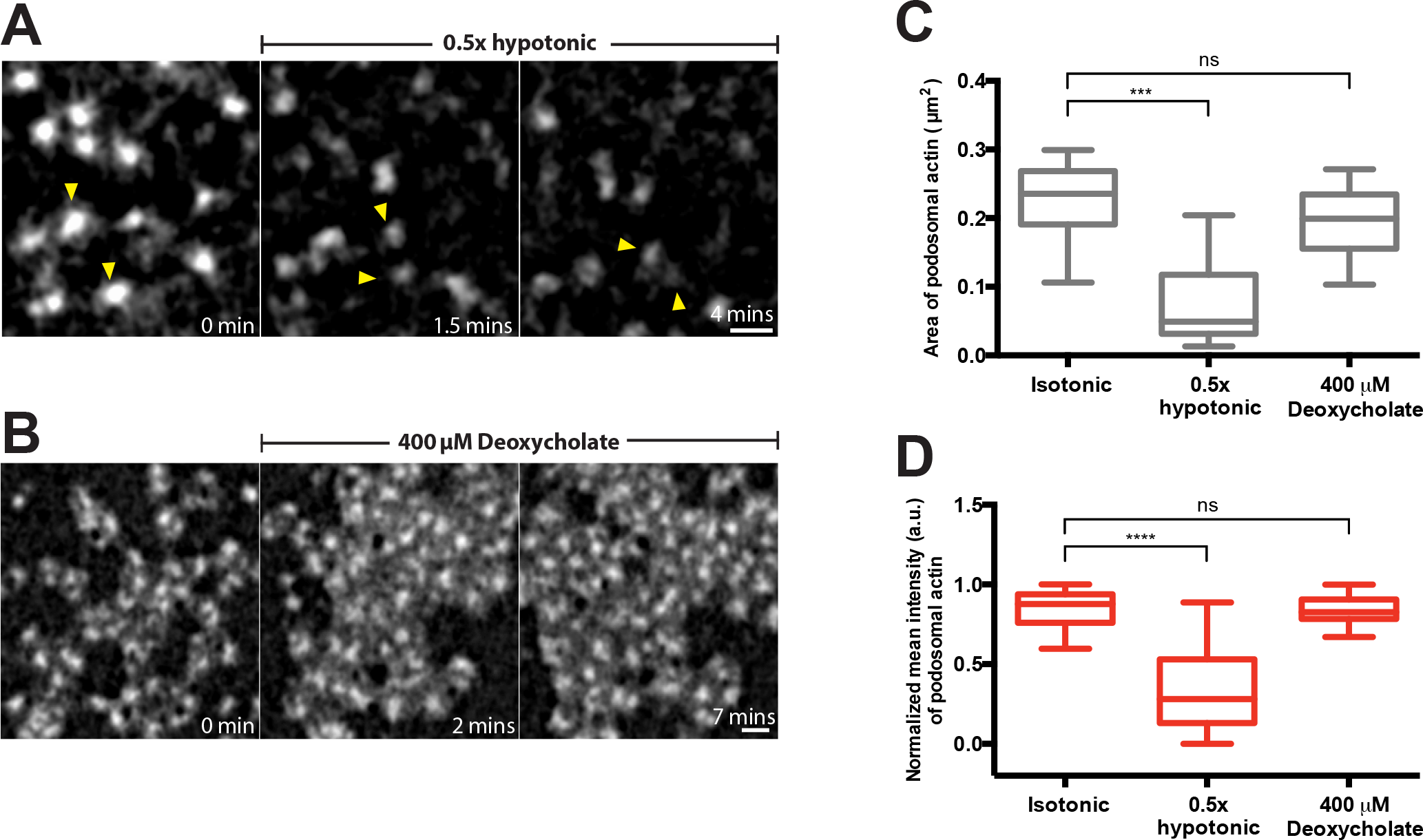
Comparison of the effects of osmotic swelling and deoxycholate treatment on podosomes. (A and B) High magnification time-lapse sequences showing evolution of podosomes in cells transfected with GFP-β-actin incubated with 0.5x hypotonic medium (A) or 400 μM deoxycholate (B). The image sequences are taken from Movie 2 and 5, respectively. Two individual podosomes in (A) are indicated by yellow arrowheads to better follow their fate. Scale bars, 1 μm. Podosome area (C) and mean actin fluorescence intensity (D) in control cells, and in cells treated with 0.5x hypotonic medium and 400 μM deoxycholate. The data are presented as a box-and-whiskers plots. At least 20 podosomes per cell were analyzed for each group. The p-values are marked by asterisks as described in the legend to Figure 1.

## Movie legends

### Supplementary Movie 1

Disruption of podosomes labeled by RFP-lifeact (left, red) in THP1 cell plated on micropatterned circular fibronectin-coated islands (4 μm diameter) upon treatment with the RhoA activator CN03 (0.1 μg/ml). Local podosome disassembly was accompanied by a massive burst of myosin-II filament assembly in each individual islands, as visualized by GFP-MRLC (right, green). Structured-illumination microscopy (SIM) was used. Single plane close to the substrate is shown. The frames were recorded at 30-second intervals over a period of 15 minutes. Display rate is 10 frames/sec. The movie corresponds to the time-lapse series shown in Figure 1D. Scale bar, 5 μm.

### Supplementary Movie 2

Dispersion and subsequent disassembly of podosomes labeled by GFP-β-actin in THP1 cell exposed to 0.5x hypotonic medium for 15 minutes. Single plane close to the substrate is shown. Structured-illumination microscopy (SIM) was used. The frames were recorded at 10-second intervals over a period of 15 minutes. Display rate is 10 frames/sec. The movie corresponds to the time-lapse series shown in Figure 2F. Scale bar, 5 μm.

### Supplementary Movie 3

Inhibition of Rho kinase by 30 μM Y-27632 in THP1 cell did not prevent the disruptive effect of 0.5x hypotonic shock on podosomes labeled by GFP-β-actin. The imaging started at the 30^th^ minute after addition of Y-27632 and was continued for another 30 minutes in the presence of both Y-27632 and 0.5x hypotonic medium. Structured-illumination microscopy (SIM) was used. Single plane close to the substrate is shown. The frames were recorded at 30-second intervals over a period of 30 minutes. Display rate is 10 frames/sec. The movie corresponds to the time-lapse series shown in Figure 2G. Scale bar, 5 μm.

### Supplementary Movie 4

Incubation of mouse fibroblast in 0.5x hypotonic medium did not affect focal adhesions visualized by mCherry-vinculin. The frames were recorded at 1-minute intervals over a period of 30 minutes using confocal microscopy. Single plane close to the substrate is shown. Display rate is 5 frames/sec. The movie corresponds to the time-lapse series shown in Figure 2H. Scale bar, 10 μm.

### Supplementary Movie 5

Addition of 400 μM deoxycholate induced transient disassembly of podosomes and subsequent formation of podosome clusters in THP1 cell. Podosomes were labeled by GFP-β-actin. Single plane close to the substrate is shown. The frames were recorded at 30-second intervals over a period of 30 minutes using SIM. Display rate is 10 frames/sec. The movie corresponds to the time-lapse series shown in Figure 3F. Scale bar, 5 μm.

### Supplementary Movie 6

The movie shows the recovery of uniform podosome distribution in cells incubated in 400 μM deoxycholate-containing medium. The cell was labeled by GFP-β-actin. The imaging started at the 30^th^ minute after addition of deoxycholate and was continued for another 30 minutes in the presence of deoxycholate. Single plane close to the substrate is shown. The frames were recorded at 15-second intervals over a period of 30 minutes using SIM. Display rate is 10 frames/sec. The movie corresponds to the time-lapse series shown in Figure 3G. Scale bar, 5 μm.

